# B-cell precursor acute lymphoblastic leukaemia with *IGH::CEBP* rearrangement: what have we learnt over the years?

**DOI:** 10.1101/2025.09.29.679048

**Authors:** A Alqahtani, KTM Fung, L Marchetti, O Heidenreich, CJ Harrison, AV Moorman, LJ Russell

## Abstract

B-cell precursor acute lymphoblastic leukaemia (BCP-ALL) is a haematologic malignancy marked by the rapid proliferation of immature B cells in the bone marrow. While BCP-ALL most commonly affects children aged 1-5 years, it remains the most prevalent subtype of ALL in adolescence and adulthood. Chromosomal translocations involving the *immunoglobulin* (*IG*) locus and partner genes are proven useful for risk stratification and guiding clinical trials for therapeutic decision. This includes translocations with *CCAAT/enhancer-binding proteins (CEBP)*, which are particularly rare. This rarity has limited efforts to characterise their genetic and clinical profiles, making risk stratification for *IGH::CEBP*-rearranged BCP-ALL challenging. In this letter, we review the clinical and demographic characteristics of all reported *IGH::CEBP* cases prior to 2024 and introduce new cases, with preliminary analysis to encourage further investigation into this poorly understood subtype. This study delivers new insights into the molecular and cytogenetic landscape of *IGH::CEBP* rearrangements in BCP-ALL, and lays a foundation for further investigation into *CEBP* family roles in haematopoietic development and leukemogenesis, especially in the context of Down syndrome. Finally, it introduces the ongoing international collaborative effort to assemble the largest known *IGH::CEBP* cohort for comprehensive risk stratification and prognostic evaluation.

---

B-cell precursor acute lymphoblastic leukaemia (BCP-ALL) is a haematologic malignancy characterised by the rapid growth of immature B cells in the bone marrow. Most BCP-ALL cases are associated with genetic abnormalities such as chromosomal translocations or point mutations^1^, which have prognostic significance. As a result, risk stratification based on chromosomal features plays a critical role in guiding patient management and informing prospective clinical trial protocols^2,3^. The identification of immunoglobulin locus (IG) translocations is proven useful for risk stratification^3^. Among these, translocations involving CCAAT enhancer binding proteins (*CEBP*) are particularly rare. The *IGH::CEBP* fusion was first reported in BCP-ALL by Chapiro et al. (2006) and Akasaka et al. (2007)^4,5^. However, the rarity of such cases has limited efforts to characterize their genetic landscape and associated clinical profiles. In this letter, we review all reported cases of *IGH::CEBP* in BCP-ALL up to 2024 to investigate potential correlations with cytogenetic aberrations, as well as clinical and demographic characteristics. Moreover, we report six new *IGH::CEBP* cases, in which we correlate their gene expression profile to the major BCP-ALL subtypes, and anticipate the interplay between CEBP family members and the affected pathways.

A total of 151 cases, including six new cases from the current study, of BCP-ALL with *IGH::CEBP* translocations have been reported in the literature prior to 2024 (Figure 1A). The majority of these cases involved *IGH::CEBPD* (n=96), followed by *IGH::CEBPA* (n=25), *IGH::CEBPE* (n=17), and *IGH::CEBPB* (n=13, Figure 1B). The white blood cell (WBC) count for these cases ranged between 1-190X10^9^/L, with median count at 16.1X10^9^/L. While the male/female ratio across all *IGH::CEBP* cases was similar (1:1.01, M:50.3%), there were more female cases associated with *IGH::CEBPA,::CEBPB*, and *::CEBPE* compared to *IGH::CEBPD* (1:1.6, M:38%, Figure 1C). The median age at diagnosis across all *IGH::CEBP* cases was 15 years (average: 19.6 years; range: 2-76 years). However, when considering the involvement of each CEBP partner gene, it was apparent that the *IGH::CEBPD* rearrangements were more prevalent in younger patients (median age: 13 years; average: 16 years; Figure 1D). This difference in age distribution was statistically significant when compared to cases involving all other *CEBP* partners (Mann– Whitney U test, p < 0.001). In contrast, individuals with *IGH::CEBPA* had a median age at diagnosis of 28 years (average: 32.2 years), those with *IGH::CEBPB* had a median of 26.3 years (average: 24.3 years), and those with *IGH::CEBPE* had a median of 16 years (average: 24.4 years; Figure 1D).

**Figure 1:**
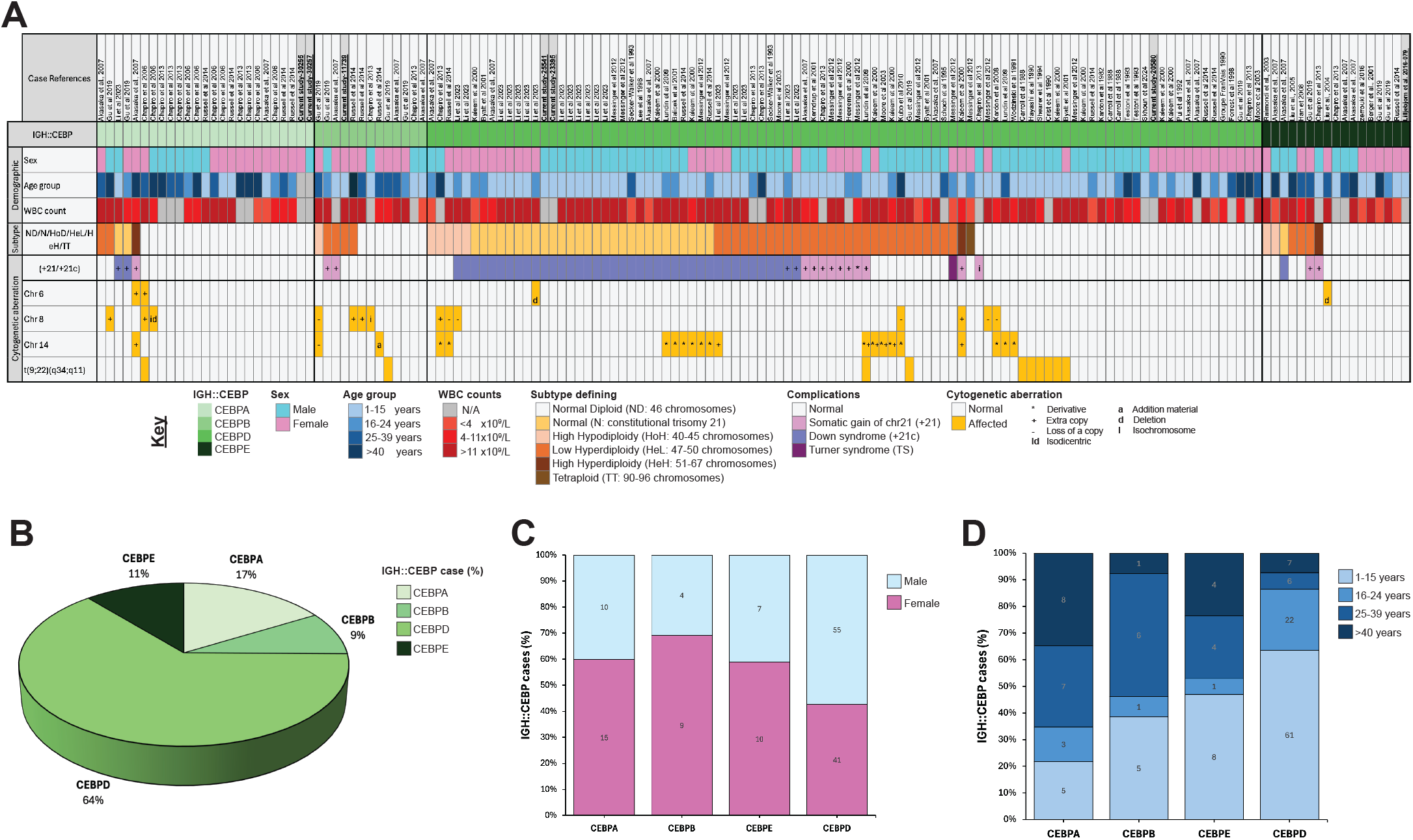
Oncoplot of *IGH::CEBP* rearranged BCP-ALL cases reported between 1982 and 2024. **A)** Clinical and cytogenetic features of the reported *IGH-CEBP* rearrangement cases in BCP-ALL (n=151). **B)** Frequencies of genetic subtypes involving an *IGH* translocation with a member of the *CEBP* gene family: CEBPA (n=25), CEBPB (n=13), CEBPD (n=96), and CEBPE (n=17). **C)** Gender distribution across *IGH::CEBP* subtypes. **D)** Age range distribution across different *IGH::CEBP* gene fusion cases.

The association of *IGH::CEBP* with Down syndrome (DS) in BCP-ALL has been previously reported and linked to poorer outcome^6^. In our analysis, 42% of cases with *IGH::CEBPD* occurred in individuals with DS; much higher than the overall rate of DS in ALL cohorts of <5%^7^. This percentage increased to 52% when including individuals with somatic gain of chromosome 21 (+21). As DS patients have a constitutional trisomy 21, the expected 47 chromosomes were observed in 73% of the cases and the remaining cases had either shown 48-49 chromosomes (22%) or 46 chromosomes (5%). The karyotype of these cases was referred to as normal (N), low hyperdiploid (HeL) and high hypodiploidy (HoH), respectively (Figure 1A). The DS-associated cases were found in individuals under 24 years of age, potentially explaining the observed enrichment of younger patients in the *IGH::CEBPD* subgroup. In contrast, translocations involving *IGH* and other *CEBP* family members showed lower prevalence in DS-ALL patients or patients with somatic gain of chromosome 21 - 12% in *IGH::CEBPA* and 18% in *IGH::CEBPE*. These findings suggest a possible unique role for *CEBPD* in the leukemogenesis of DS-associated BCP-ALL. Further investigation is required to elucidate the mechanisms by which *CEBPD* contributes to leukemic transformation in the context of DS.

Cytogenetic aberrations, such as trisomy 6 or deletions of chromosome 6, have been observed in a subset of ALL cases^8,9^. In the cohort of cases with *IGH::CEBPA*, we identified two instances with trisomy 6 and two additional instances with chromosome 6 deletion, distributed among *IGH::CEBPD and::CEBPE* cases (Figure 1A). The gain of chromosome 6 in one *IGH::CEBPA* case was associated with high hyperdiploid (HeH) karyotype, a feature that has been linked to favourable prognosis in ALL^8^. However, the deletions of 6q21, which includes several known tumour-suppressor genes, could contributed to malignant transformation or proliferation^9^. In addition, chromosome 8 abnormalities were identified in 14 *IGH::CEBP* cases. Of these, 6 showed gain of chromosome 8 (in which two were associated with HeH, another two with HeL, one with HoH and one with normal diploidy), another 6 had complete deletion of chromosome 8, and 2 exhibited either an isochromosome i(8)(q10) or isodicentric chromosome idic(8)(p11). These alterations were distributed across *IGH::CEBPA* (n=3), *:: CEBPB* (n=4), and *::CEBPD* (n=7) cases (Figure 1A). Many of these cases (n=10/14) occurred in adults, similar to the previously reported cases of chromosome 8q24 aberrations being more frequent in adult ALL^10^. Somatic gain of chromosome 14 or trisomy 14 is relatively common in hyperdiploid BCP-ALL. In our cohort, we noted trisomy 14 presence in 7 cases, primarily associated with *IGH::CEBPD* and occurring in the context of a HeL karyotype (n=6/7, Figure1A). Moreover, in 16 *IGH::CEBPD* cases, the translocation between chromosomes 14 and *CEBPD* resulted in a derivative chromosome 14. This indicates that the centromere of chromosome 14 was retained, placing *CEBPD* under the regulatory control of the *IGH* super-enhancers, and leading to overexpression of *CEBPD* (Figure 1A). These findings highlight the need for further investigation to elucidate the biological and clinical implications of these chromosomal alterations in *IGH::CEBP* driven BCP-ALL.

The Philadelphia chromosome, resulting from the reciprocal translocation, t(9;22)(q34;q11), and generating the *BCR::ABL1* fusion, was identified in 7% of all reported *IGH::CEBP* cases (n=10/151). This was most prominent in *IGH::CEBPD*, where it occurred in 8% of cases (n=8/96, Figure 1A). Interestingly, all these cases were exclusive from DS and in patients under the age of 15 years old, which appeared to be higher than the reported percentage (3-5%) for *BCR::ABL1* fusion in paediatric ALL^11^. The coexistence of *BCR::ABL1* and *IGH::CEBP* rearrangements remains largely unknown and may have significant implications for both disease pathogenesis and prognosis. The *BCR::ABL1* fusion leads to persistently enhanced tyrosine kinase activity, whereas *CEBP* gene fusions result in aberrant overexpression of transcription factors critical for hematopoietic differentiation. Together, these alterations may synergistically disrupt normal B-cell development and function, contributing to the initiation and progression of BCP-ALL.

In this preliminary study, we analysed the transcriptome of 7 cases with *IGH::CEBP* rearrangements from a larger cohort of 188 BCP-ALL cases, including 182 cases from EGAS00001001795^12^ and 6 new cases we collected (Figure 1A). RNA sequencing was performed on diagnostic samples, and molecular subtyping was conducted based on gene expression profiling with reference to Li et al. (2018)^13^. In total, we identified eight distinct molecular subtypes of BCP-ALL, visualized using uniform manifold approximation and projection (UMAP; Figure 2A). The *IGH::CEBP* cases were distributed across two clusters, including one associated with hyperdiploid expression profile. The hyperdiploid cluster included cases with HeL, HeH, and TT karyotypes. To further explore molecular changes among the *IGH::CEBP* cases, we performed hierarchical clustering based on the top 300 differentially expressed genes (DEGs). This analysis revealed that the five cases of *IGH::CEBPB* (n=1), *::CEBPD* (n=3), and *::CEBPE* (n=1), exhibited a similar gene expression signature. In contrast, the two *IGH::CEBPA* cases formed a separate cluster, indicating a distinct transcriptional profile (Figure 2B). Furthermore, KEGG pathway analysis of the top 300 DEGs identified significant enrichment in pathways related to cancer and transcriptional deregulation in cancer (Figure 2C). In the three cases of *IGH::CEBPD*, there were two cases with DS (Asterisk in Figure 2B). The hierarchical clustering indicated that the two DS-associated *IGH::CEBPD* cases shared a similar gene expression signature, while the non-DS case showed greater similarity to the *IGH::CEBPE* case (Figure 2B). Moreover, the two DS-associated *IGH::CEBPD* cases exhibited higher *CEBPD* expression levels compared to the third case without DS (asterisk in Figure 2D). Overexpression of *CEPBD* in a DS mouse model was reported to alter the genetic context of B cell development, resulting in a persistent predominance of pro-B cells. In contrast, *CEBPD* overexpression in wild-type controls promoted increased B-lineage differentiation^6^. Hence, these findings could suggest that the combination of *CEBPD* overexpression and trisomy 21 may contribute to distinct gene expression patterns in DS-associated *IGH::CEBPD*, potentially increasing susceptibility to BCP-ALL. Further investigation is needed to elucidate the biological mechanisms underlying this interaction in human cells and to determine its potential impact on dysregulating B cell development in BCP-ALL.

**Figure 2:**
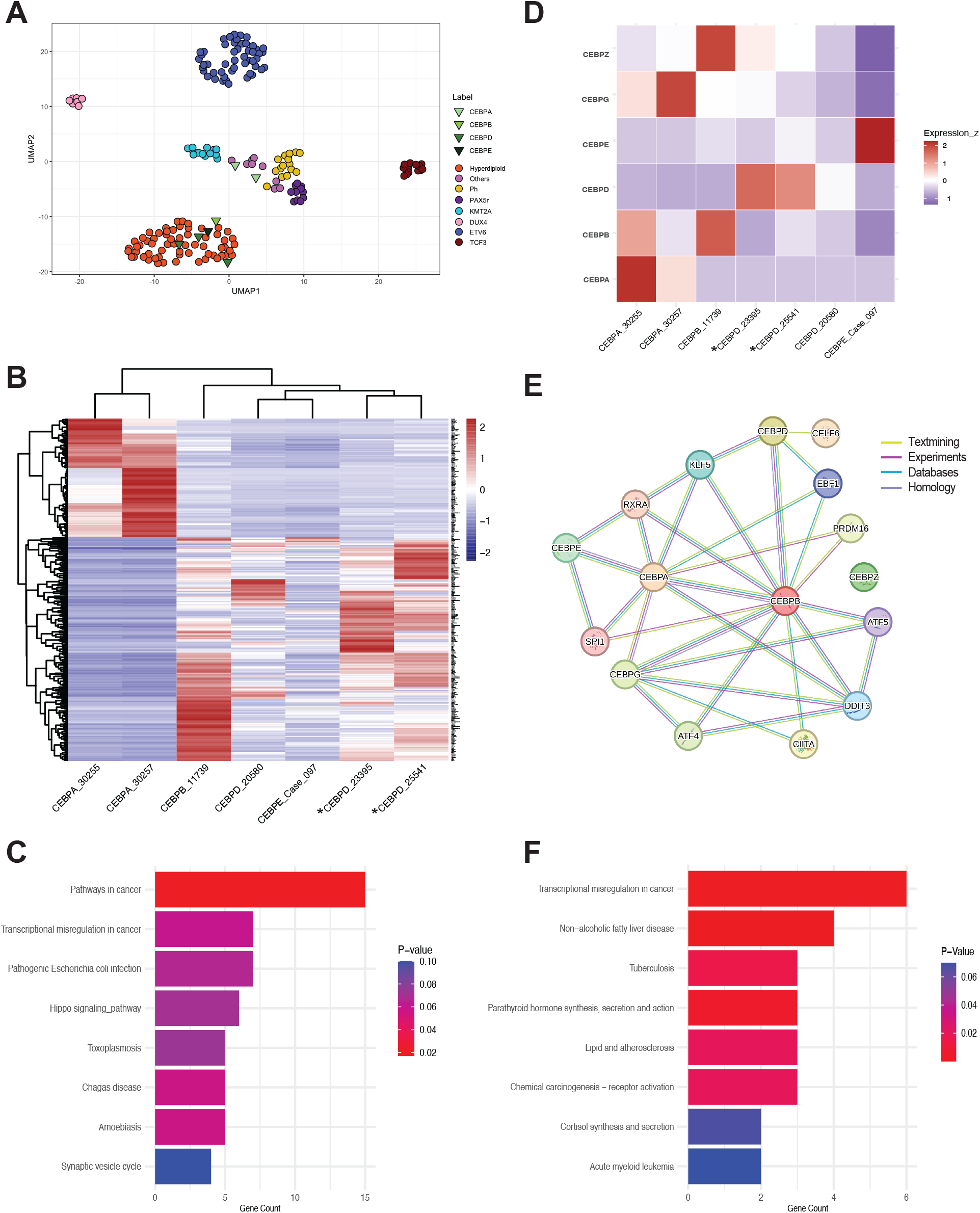
Transcriptomic and genomics of *IGH::CEBP* cases. **A)** Gene expression of 188 BCP-ALL cases visualized by UMAP, showing the *IGH::CEBP* cases (n=7, Green) distributed across different clusters. *IGH::CEBPA* cases were clustered separately from the other *IGH::CEBP* cases. The average silhouette score was used to generate this 2-dimensional UMAP. Clusters and CEBP cases are shown in different colours. **B)** Hierarchical clustering of the *IGH::CEBP* cases identified that the two *IGH::CEBPA* cases had a unique gene expression signature (padj > 0.01) compared to the other *IGH::CEBP* cases. Remarkably, the *IGH::CEBPD* cases with DS clustered together, separate from the *IGH::CEBPD* without DS. **C)** KEGG pathway analysis of the top 300 DEGs in *IGH::CEBP* cases showed enrichment for pathways involved in cancer and transcriptional dysregulation. **D)** Heatmap presenting differences in gene expression between *IGH::CEBP* cases. Interestingly, *IGH::CEBPD* cases with DS showed high expression of *CEBPD* compared to the *IGH::CEBPD* case without DS. Moreover, expression of *CEBPG*, which is involved in the tight regulation of B cell development, was high in the two *IGH::CEBPA* cases and this can potentially contribute to their distinct expression profile (see panel B). **E)** Protein–protein interaction analysis using the STRING database was performed to investigate the functional interactions among CEBP family members and their associated proteins. A maximum of 10 interactors were included, filtered by a high-confidence interaction score (>0.7). Yellow: Textmining, purple: experiments, blue: database, light purple: homology. **F)** KEGG pathway analysis of CEBP family members and their associated proteins revealed enrichment in pathways related to transcriptional dysregulation and acute myeloid leukaemia.

B cell development involves tight regulation of *CEBP* family members^14,15^. While CEBPγ and possibly CEBPζ can act as negative regulators of other CEBP proteins— modulating their transcriptional activity through inhibitory interactions^15,16^— their roles in B cell development or BCP-ALL remain largely unknown. Therefore, we examined the expression of these negative regulators in *IGH::CEBP* cases to determine whether they might influence the unique genomic features observed in these cases. Remarkably, *CEBPG* was specifically overexpressed in *IGH::CEBPA* cases (n=2), while increased expression of *CEBPZ* was observed in both *IGH::CEBPB* (n=1) and DS-associated *CEBPD* cases (n=2/3, Figure 2D). This hence may suggest a role for these inhibitors in the modulating other *CEBP* family member activities.

The functional protein associations of CEBP family members were analysed using the STRING database (https://string-db.org). This analysis revealed that CEBPα and CEBPβ were central nodes in the network, associated with each other and with other CEBP family members—including CEBPγ (for both), CEBPε (for CEBPα), and CEBPδ (for CEBPβ). However, CEBPδ and CEBPε showed no interactions with each other or with CEBPγ (Figure 2E). We included the top ten interacting proteins with high-confidence interaction scores (≥0.7), which encompassed factors involved in lymphoid and myeloid lineage development, such as SPI1, ATF4/5, and DDIT3 (Figure 2E). This network was most significantly enriched for KEGG pathways related to transcriptional dysregulation in cancer, carcinogenesis, and acute myeloid leukaemia—consistent with the pathway enrichment observed in the *IGH::CEBP* cases (Figure 2F). Therefore, the interaction profile of CEBP family members highlights their central role in transcriptional regulation pathways implicated in haematopoietic development and leukemic transformation.

Altogether, this study provides new insights into the molecular and cytogenetic landscape of *IGH::CEBP* rearrangements in BCP-ALL. Although most reported cases lack comprehensive treatment information and outcome data, making it difficult to assess prognostic implications and inform risk stratification, we are conducting an international study to assemble a larger cohort of *IGH::CEBP* cases to address these gaps. In addition, our findings highlight the clinical and biological heterogeneity of *IGH::CEBP* fusions and emphasise on the need for further investigation into their role in B-cell development, particularly in the context of DS and co-occurring genetic alterations.

## Acknowledgements

We would like to thank the Blood cancer UK (BCUK, 24012) for funding this project.

